# Comparing average network signals and neural mass signals in systems with low-synchrony

**DOI:** 10.1101/196113

**Authors:** P. Tewarie, A. Daffertshofer, B.W. van Dijk

## Abstract

Neural mass models are accepted as efficient modelling techniques to model empirical observations such as disturbed oscillations or neuronal synchronization. Neural mass models are based on the mean-field assumption, i.e. they capture the mean-activity of a neuronal population. However, it is unclear if neural mass models still describe the mean activity of a neuronal population when the underlying neural network topology is not homogenous. Here, we test whether the mean activity of a neuronal population can be described by neural mass models when there is neuronal loss and when the connections in the network become sparse. To this end, we derive two neural mass models from a conductance based leaky integrate-and-firing (LIF) model. We then compared the power spectral densities of the mean activity of a network of inhibitory and excitatory LIF neurons with that of neural mass models by computing the Kolmogorov-Smirnov test statistic. Firstly, we found that when the number of neurons in a fully connected LIF-network is larger than 300, the neural mass model is a good description of the mean activity. Secondly, if the connection density in the LIF-network does not exceed a crtical value, this leads to desynchronization of neurons within the LIF-network and to failure of neural mass description. Therefore we conclude that neural mass models can be used for analysing empirical observations if the neuronal network of interest is large enough and when neurons in this system synchronize.

## 2 Introduction

There is an increasing need in neuroscience to comprehend the enormous amount of empirical data obtained with neuroimaging modalities such as fMRI and MEG/EEG by means of theoretical analysis. Computational approaches such as neural mass models are widely used in this context (Deco et al., 2008). The advantage of neural mass models is that they simulate the average activity of all neurons in a neuronal population, and thereby avoid the overwhelming detailed behaviour of all individual neurons in a neuronal population. However, neural mass models are based on a mean-field approximation, which assumes homogeneity of the underlying neuronal population. As realistic neuronal populations are naturally inhomogeneous, it is unclear to what extent neural mass models are able to capture the average activity of these realistic populations.

The mean field approximation has its roots in statistical physics where it is often applied to reduce many body systems to low dimensional systems (Landau and Lifshitz, 1968). For instance, mean-field theory has been used in ising spin systems to replace individual interactions between atomic spins by an effective mean-field which accounts for the average interaction over all local atomic spins. However, it is known from these systems that a mean-field description fails when this average interaction term is dependent on spatial location (e.g. inhomogeneity), when fluctuations around this average are large or when the size of the system becomes too small.

Mean-field theory has been widely applied in neuroscience where the underlying assumption is it that each neuron in a neuronal population can be described by equivalent statistics and that each neuron converges to the same attractor in phase space (Deco et al., 2008). This allows to describe the dynamics of neuronal population by a limited number of time dependent variables corresponding to the average dynamics of the population (Baladron et al., 2012). The very first studies in neuroscience demonstrated analytically that the mean-field approximation was valid for homogenous neural networks when the numbers of neurons tend to infinity (Amari, 1972; Amari et al., 1977). Ever since, numerous studies have analysed the mean-field in neural networks under diverse conditions such as in finite neural networks (Mattia and Del Giduice, 2002; Touboul and Ermentrout, 2011), with different connectivity patterns (Cessac and Vieville, 2013; Moynot and Samuelides, 2002; Samuelides and Cessac, 2007; Brunel and hakim, 1999; Cessac, 1995), using different single-cell neuronal models (Abbot and Vreeswijk, 1993; Cessac, 2008; Treves, 1993)), using realistic parameter regimes (Grabska-Barwinska and Latham, 2013), or as a function of time (Molgedey et al, 2013; Moynot and Samuelides, 2002), the so-called dynamic mean-field approaches.

Despite the large number of previous studies on mean-field approaches in neural networks, both the derivation of neural mass models from single-cell neuronal models and the effect of mean-field approximations involved in this derivation to describe the average activity of realistic inhomogeneous neuronal populations are not fully elucidated (Breakspear et al., 2003; Faugeras et al., 2009; Wong and Wang, 2006). In notable work by Rodrigues and colleagues (2010), two mappings between conductance based leaky integrate-and-fire (LIF) neurons and a commonly used neural mass model were introduced that allowed for an interpretation of model characteristics between the two scales (Rodrigues et al., 2010). The two models were based on different assumptions with regard to the time scales of the membrane and synaptic activities of neurons, as these time scales were either considered similar or dissimilar. Irrespective of the assumption related to these time-scales, both models resulted in a neural mass model similar to that of the well-known Freeman model (Freeman, 1975). Although these findings form a basis for future research, neurons in the simulated neuronal populations in this study were kept in a non-spiking regime which is not a realistic representation of the underlying physiology as spiking is a prerequisite for neuronal populations to communicate between each other.

In the present study we build upon the work of this previous study and compare neural mass signals with the average signal of inhomogeneous neuronal populations in which individual neurons operate in the spiking regime (Rodrigues et al., 2010). Our main objective is to analyse what the effect of mean-field approximations are for neural mass models when the underlying neuronal population is inhomogeneous. In addition, we investigate the lower limit of neuronal network size for which a neural mass description is still valid. In this study, we use a network of conductance based inhibitory and excitatory LIF neurons to model a neuronal population and we derive two neural mass models from the LIF model. Since it is extremely challenging to solve this comparison problem analytically, we adopt a numerical approach to compare average signals from neuronal populations with neural mass signals. During these analyses, we either decrease the number of neurons or decrease the number of connections in the neuronal population for different network topologies. Lastly, we hypothesize that a discrepancy between the average signal of a neuronal population and the neural mass signal can potentially be explained by desynchronization between neurons in the neuronal population.

## 3 Neuronal models

### 3.1 Network of leaky integrate-and-fire neurons

We consider a population of *N* LIF neurons (Rudolph-Lilith et al., 2012). A LIF neuron can be regarded as a parallel electric circuit with a resistor, capacitor *C* and an external input current *J*_*ext*_. Every neuron is described in terms of the dynamics of its membrane potential *V*_*j*_ with *j* = 1*,…, N* and its time-dependent synaptic conductances. Conductances are denoted as 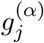 for every neuron *j*, which alter due to incoming spikes. Synapses are either excitatory or inhibitory and can be discriminated by *α*, where *α ∈* {*E, I*}. We assume that *V*_*j*_ obeys the stochastic dynamics

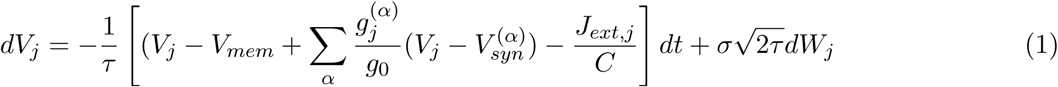

with fixed resting state potential *Vmem* and reversal potentials *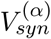 g*_0_is the leak conductance that is considered constant and *τ* is the membrane time constant. Both constants are assumed to be identical for all neurons. The membrane time constant *τ* is related by 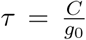 to the capacitor and membrane conductance constant *g*_0_. *τ* determines the rise and decay of the membrane potential and *J*_*ext,j*_ refers to an external current on neuron *j*. Every neuron j is under influence of zero-centered, *δ* correlated, Gaussian white noise *W*_*j*_ with unit variance, i.e. that is 𝔼 [*W*_*j*_ = 0 and 𝔼 [*W*_*j*_(*t*)*W*_*j*_(*t′*)] = *δ_jk_δ*(*t – t′*) where *δ_kl_* is the Kroneckers delta function and *δ*(*t - t′*) refers to Diracs delta-function. The noise strength *σ* is considered identical for all neurons *-* see Table 1 for values of all constants (Vogels and Abbott, 2005). To simplify our approach and to increase the interpretability of our analyses with previous LIF studies we assume that conductances have an infinitely fast rising time (Yger et al., 2011). This can be assumed if neurons of interest operate in the high frequency regime which is usually the case for excitatory neurons and often also for inhibitory neurons (Creutzfeldt and Ito, 1968). Therefore, the dynamics of the conductances can be described by a first order response and depend on the incoming spikes as follows

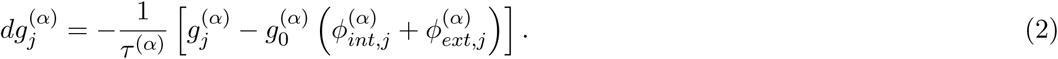

**Table 1:** Constants used in simulations.

|  |  |  |
| --- | --- | --- |
| $\tau$ | Membrane time constant | 20 ms |
| $V_{mem}$ | Resting state potential | -60 mV |
| $V_{syn}^{(E)}$ | Reversal potential for excitatory synapses | 0 mV |
| $V_{syn}^{(I)}$ | Reversal potential for inhibitory synapses | -80 mV |
| $V_{thres}$ | Threshold potential | -50 mV |
| $V_{reset}$ | Reset potential | -60 mV |
| $g_0$ | Leak conductance | 10 nS |
| $g_0^E$ | Excitatory synaptic conductance | 3 nS |
| $g_0^I$ | Inhibitory synaptic conductance | 50 nS |
| $\Delta t_{refract}$ | Refractory period | 5 ms |
| $\sigma$ | Noise strength | 0.6 mV |
| $\tau^{(E)}$ | Excitatory synaptic time constant | 5 ms |
| $\tau^{(I)}$ | Inhibitory synaptic time constant | 10 ms |

Here 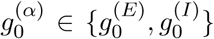 refer to constants related to the maximal conductance that serves to discriminate between excitatory and inhibitory neurons. *τ* ^(*α*)^ determines the decay of synaptic activity which is in general different for inhibitory and excitatory neurons. 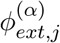 and 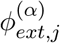 corresponds to incoming firing rates from respectively internal and external sources. Due to variation in synaptic conductances the membrane potential of neurons will fluctuate. If these membrane potential fluctuations reach the threshold potential, the neuron spikes and the membrane potential is set to a reset potential and kept in a short refractory state of 5 ms.

### 3.2 Neural mass models

We derive two neural mass models from the LIF neuronal model. See appendix B for a detailed derivation of both neural mass models. Here, we briefly mention the assumptions and subsequent approximations that are necessary to obtain both neural mass models.

For the first mapping we follow an approach introduced by Rodrgues and colleagues (2010). Based on some experimental indications, the starting assumption here is that we assume that membrane and synaptic time scales are of the same order of magnitude (Destexhe and Sejnowski, 2001; Gerstner and Kistler, 2002). We consider the expectation value of the membrane potential *V* (from Eq. 1), main variable of interest, and the expectation values of the synaptic conductances *g*^(*α*)^ and firing rates *φ_int_* for inhibitory and excitatory neurons separately (from Eq. 2). Subsequently, we assume that fluctuations of driving forces 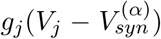 for synaptic currents (in Eq. 1) are relatively small which allows for a mean-field approximation 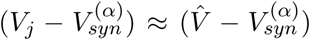 which simplifies the expression for *V*. Then, the next assumption entails that time scales of synaptic activity are of the same order irrespective their type, which allows us to combine excitatory and inhibitory synaptic activity into one description. These assumptions lead to modified first order differential equations (for Eqs. 1 and 2 describing *V* and *g*) and combining these yield a second order differential equation as description for our first neural mass model (Freeman, 1975)

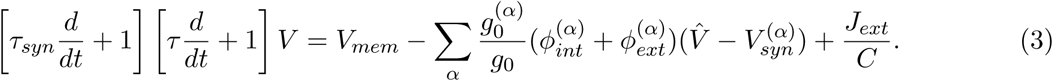

Note that the index *j* is omitted in the equation as variable *V* does not corresponds to single neurons anymore but to the expectation value of *V*_*j*_ and hence the neural mass. We further ignored the reset-rule and the presence of a refractory period, incorporated in the initial LIF equations, as we assume that the most important characteristics that need to be captured are sub-threshold post-synaptic potentials of neurons in the neuronal population. Notice that our first neural mapping model is identical to the widely known neural mass model of Freeman (see Appendix B) (Freeman, 1975). 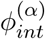 and 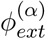 in Eq. 3 refer to average firing incoming to neurons in the population which are assumed to be deterministic and defined as

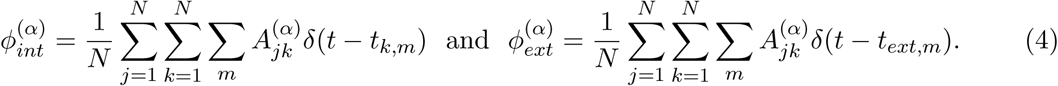

Here, *δt - t*^*I*^ refers to the Diracs delta-function. *t*_*k,m*_ is the time of the *m*-th spike of neuron *k* and synapse *j, k, α* may also receive external input in the form of spikes at times 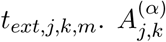 is the adjacency matrix describing whether neuron *k* targets neuron *j*.

For the second mapping we question some of the key assumptions made in the first mapping and therefore follow a different route. First of all, we question the assumption that driving currents should be considered as constants. Related to this, it is not entirely clear in the former approach why this assumption only applies to synaptic driving currents but not to the leaky driving current. In the second neural mass mapping we follow neural-field like approach. Here, we again consider the expectation values of *V*_*j*_, *g* ^(*α*)^, 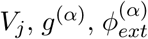 and 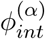 and similarly assume that time scales of synapses agree irrespective of their type. We now combine both first order differential equations of *V* and *g*^<^, which results in a second order differential equation. The main approximation related to neural field-theory is that we follow the assumption that the time scale of the membrane potential is much smaller than the time scale of the synaptic conductivity. This indicates that the membrane potential instantly follows changes at the synapse. Therefore, the dynamics of the membrane potential can be eliminated adiabatically which eventually yields for the mean membrane potential of the second neural mass model

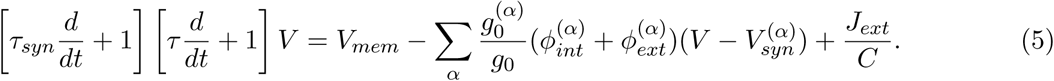

Qualitative difference to Eq. 3 is that Eq. 5 contains an extra parametric force term 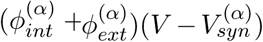 on the RHS due to which this no longer agrees with Freemans model. We again assume that 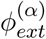 and 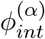 are considered to be deterministic and are defined similarly as in Eq. 4.

## 4 The LIF-network vs neural mass models

### 4.1 Simulation of average LIF-network signals

Simulation of a neuronal population of *N* LIF neurons was performed using Eqs. 1, 2, 12 and 15 with parameters defined as in table 1. Eqs. 12 and 15 can be found in the appendix and describe the incoming firing rates to each neuron, and the reset and refractory rule, respectively. See appendix C for the algorithm that was used for this purpose. For all simulations we used a ratio of 5:1 for excitatory versus inhibitory neurons (Yger et al., 2011). We consider a 1D network topology where connections between neurons were established using an adjacency matrix which is based on the network configuration of interest (see below). Simulations were executed for 50 seconds with an integration time step of 0.1ms (i.e. a sample frequency of 10 kHz). We used an initial transient current to all neurons of *J*_*ext,j*_ = 20*nA* during the first 20ms of the simulations to trigger network activity. We choose not add a constant external current input *J*_*ext,j*_ or an external firing input 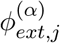 during the whole simulation time since such an external input would refer to neuronal influence from other populations or experimental stimuli which needed to be excluded. Subsequently, we used an Euler forward method to solve the coupled first order differential equations for synaptic conductances *g*_*j*_ and an Euler forward method with additive white noise in the Stratonovich picture to solve the stochastic differential equations for the individual membrane potentials *V*_*j*_. At the end of the simulations we stored all individual spike times *t*_*j,m*_ to compute the average firing and subsequently computed the average network activity 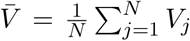 The average network activity was main variable of interest for the LIF populations and their average firing rates were stored and used in subsequent neural mass simulations.

We compared average network activity with our neural mass mapping for different network conditions:

1. Effect of network size: We ran simulations for different network sizes given a fully connected network configuration. Simulations were executed in steps of Δ*N* = 100 for 100 *< N <* 1000 and in steps of Δ*N* = 500 for 1000 *< N <* 10000.
2. Effect of network density: We ran simulations for three different network topologies: regular networks, small-world networks and random networks for networks with a size of *N* = 10000. Regular networks were defined as networks were the numnber of connections was equal for every node. Small-world networks were obtained by randomly rewiring a regular network with a probability of 0.1 and random networks were obtained by simulating Erdös R´enyi networks (Watts and Strogatz, 1998; Erdös R´enyi, 1965). For each network configuration we ran simulations for different network densities between 0 *< density <* 1.0. Networks were reconstructed using the contest toolbox for Matlab *(http://www.mathstat.strath.ac.uk/outreach/contest/)*.

### 4.2 Simulation of neural mass signals

We simulated neural mass signals by solving Eqs 3 (neural mass model 1) and 5 (neural mass model 2) by using a conventional Euler forward method. To this end, parameters in table 1 were used and the synaptic time constant in the LHS of these equations was defined as *τ_syn_* = (*τ* ^(*E*)^ + *τ* ^(*I*)^)*/*2. We used 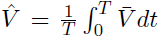 for the mean potentials temporal average in the RHS of Eq. 3. The input to both neural mass models was achieved by using the average firing rates of the simulated LIF population in the RHS of Eqs 3 and 5 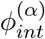 Similar to the simulations of the LIF-network activity we ignored external firing input 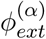 and the external input current *J*_*ext*_. We again simulated 50 seconds of data for both neural mass models with an integration time step of 0.1ms.

### 4.3 Comparing average LIF-network signals and neural mass signals

The neural mass signal *V* and the average network signal *V* were compared by testing whether the spectral contents of the signals differed. To this end, we compared the power spectral densities of the two signals by means of the two-sided Kolmogorov-Smirnov (KS) test. The two sided KS-test quantifies the distance between two empirical distibution functions and is sensitive for differences in location and in the shape of the distribution functions. In order to compare both signals, both signals were filtered using a Chebyshev low pass filter with a stop frequency of 80Hz and a pass frequency of 70Hz. Subsequently, both signals were z-transformed to ensure that the mean was zero-centered and sealing to unit variance. Power spectral densities were computed using the Welch’s periodogram method with a window of 3 seconds and an overlap of 0.4. See appendix C for the pseudo-code that was used to compare average network signals with neural mass signals.

### 4.4 Synchronization properties of the LIF-network

We computed the phase locking value (PLV) between neurons in the LIF-networks to test whether a discrepancy between average network signals and the neural mass signals could be explained by desynchronization between neurons in the LIF-network (Mardia, 1972). For this purpose we filtered the individual membrane potentials of neurons in the LIF-network with a Chebyshev band-pass filter between 8-13 Hz. Subsequently, we computed the analytical signal *a*(*t*) to extract the phase *ϕ* of the signals

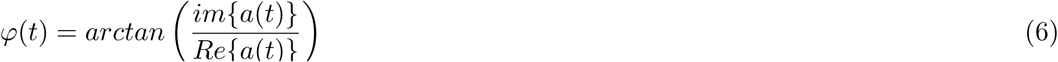

The PLV was then computed between neurons *j* and *k* in time interval *T* to determine the amount of phase synchronization between signals of individual neurons

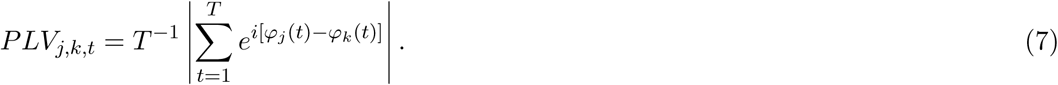

The PLV value between 1000 random pairs of neurons in the network was calculated. The mean PLV and standard error was then computed and used for further analyses. All analyses were executed using Matlab R2011b (Natick, Massachusetts: The MathWorks Inc., 2011).

## 5 Results

### 5.1 Effect of network size

We simulated fully connected LIF-networks for a range of network sizes. For the first neural mass model, we observed that there was discrepancy between the average network signal and the neural mass signal when the number of neurons was less than 300 (Fig. 1C), i.e. in the lower limit. Below the limit of 300 neurons, the Kolmogorov-Smirnov test statistic had a significant p-value (p-value *<*0.05), indicating a difference between power spectral densities of the average network signal and the neural mass signal. In Fig. 1E we show the power spectral densities with confidence intervals belonging the neural mass signal (dashed lines) and the average network signal (normal lines) for a LIF-network with 100 neurons. There is still an overlap between the two in the higher alpha band and beta band regime but it can be observed that the neural mass model failed for especially higher frequencies. For a LIF network of 2000 neurons it can be observed that there was a large overlap between the confidence intervals of the power spectral densities of the average network signal and the neural mass signal for a wide range of frequencies (Fig. 1F). Only for the highest frequencies we observe a slight shift between the two. For the second neural mass model, we again observe that there was a discrepancy between the average network signal and the neural mass signal when the number of neurons was less than 300 (Fig 1D). Again we plotted the power spectral densities with confidence intervals below the limit of 300 neurons and observe a discrepancy between the average network signal and the neural mass signal for a large amount of the frequency range (Fig 1G.). There was overlap only in the alpha frequency range. For a LIF network of 2000 neurons we again observed a large overlap between the confidence intervals, except for the highest frequencies.

**Figure 1:**
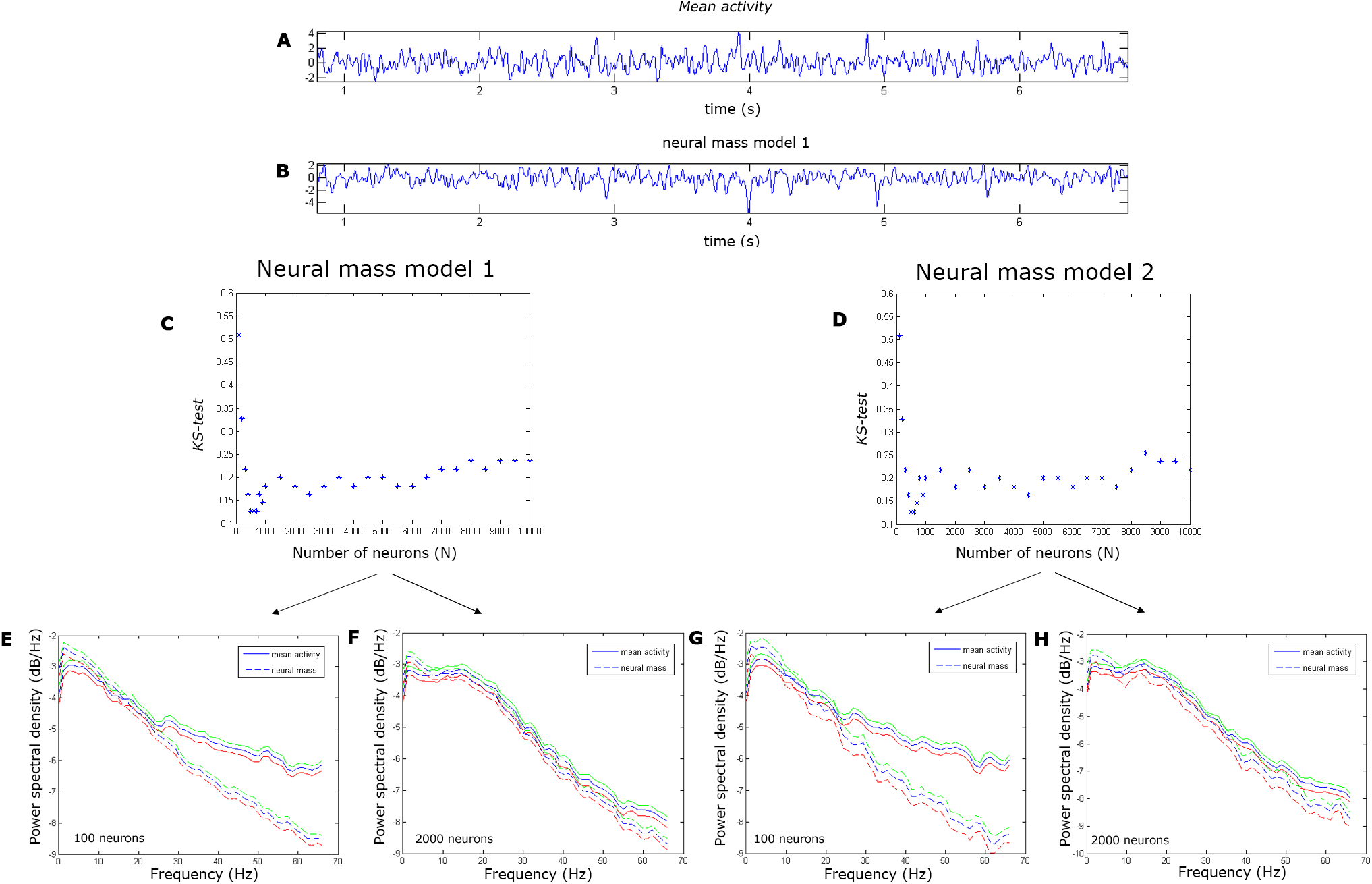
Effect of network size. Simulations were done for fully connected LIF-networks, starting with 10000 neurons down to 100 neurons. For each network size we estimated the average network signal, example shown for 1000 neurons (A). We calculated neural mass activity for both neural mass models with the mean spike train of the LIF-network as input, example shown for neural mass model 1 with 1000 neurons (B). The Kolmogorov-test statistic was used to compare the power spectral densities of the average network signals with signals from neural mass model 1 (C) and neural mass model 2 (D) for networks with increasing number of neurons. We further show conditions where neural mass models are capable (F and H) and not capable (E and G) of describing the average network signal. In each of these plots the power spectral densities with confidence intervals (red and green lines) of the average network signal (normal lines) and the neural mass signals (dashed lines) are plotted for a given frequency range, for 100 neurons (F and H) and 2000 neurons (E and G). Figure E and F correspond to neural mass model 1 and G and H to neural mass model 2.

For both neural mass models we observe that the Kolmogorov-Smirnov test statistic has local minima for networks between 500 and 1000 neurons. For larger network sizes, we observed that the Kolmogorov-Smirnov test statistic increased as a function of number of neurons, though not reaching significance.

### 5.2 Effect of network density

For three different network configurations (regular, small world and random network topology) we assessed the effect of decreasing the density of connections in the LIF-network on the ability of the neural mass model to describe the average network signal. Since we obtained similar results for neural mass model 1 and neural mass model 2 in the previous analyses, we continued these analyses by using neural mass model 1 only due to less computational expense of neural mass model 1.

For the LIF-network with a regular network topology we observed that there was a significant difference between the average network signal and the neural mass signal when the density of connections in the LIF-network was smaller than 0.9 (Fig. 2A). For LIF-networks with small world and random network topology we also observed a critical density below which there is a discrepancy between the average network signal and the neural mass model. These critical densities were 0.85 and 0.05 for the small world and random LIF networks respectively (Fig 2B & 2C). By analyzing the PLV in the LIF-networks as a function of connection density we also observed that there was an instantaneous shift to larger PLVs if the densities were higher than the aforementioned critical densities. Note that the increase in PLV beyond these thresholds was relatively small (in order of 0.1-0.2), but appeared to be instantaneous. When inspecting the power density spectra of the average network signal and neural mass signal we observed a large overlap between the confidence intervals of the average network signal and neural mass signal after the critical densities for any network configuration (Fig. 3BDF). Before the critical densities we again observed a small overlap especially in the alpha band for regular and random networks (Fig. 3AE). However, for small world LIF-networks we found that there can be still overlap between the confidence intervals up to frequencies in the beta band (Fig. 3C). For higher frequencies we again observed that the confidence intervals diverge. For all network configurations, there is higher prominence of higher frequencies in the average network signal which is not captured by the neural mass signal.

**Figure 2:**
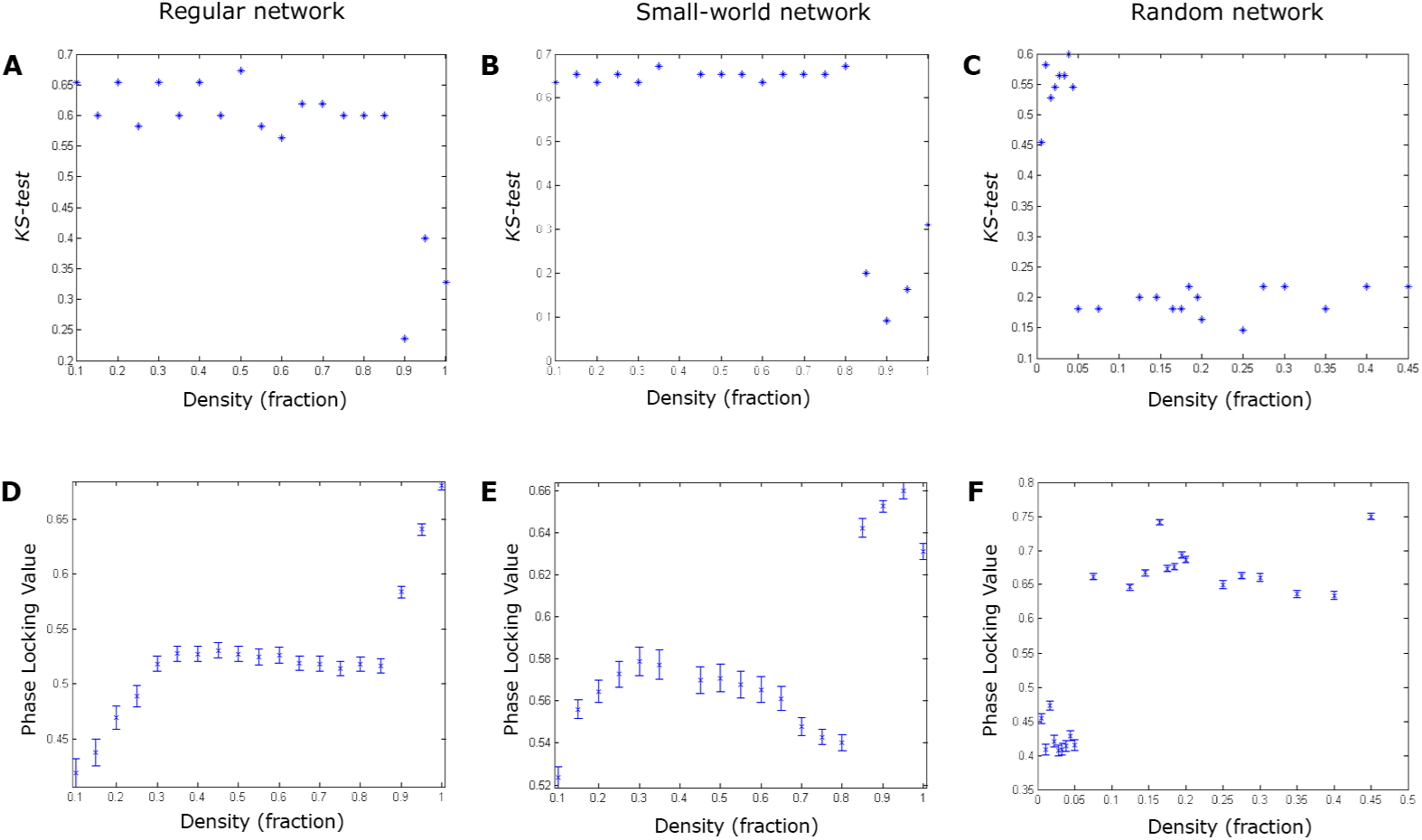
Effect of network density. Simulations were done for regular, small-world and random networks given a network size of 10000 neurons. For each network configuration we decreased the density of connections and the Kolomogorov-Smirnov test statistic was used to compare the power spectral densities of the average network signals with the first neural mass model (ABC). For each network configuration we computed the mean PLV between neurons in the neuronal network. We plotted this mean PLV (with standard error) as a function of density (DEF) for each network configuration. Note the critical thresholds for density for mean PLV values and the Kolomogorov-Smirnov test statistic for each network configuration.

**Figure 3:**
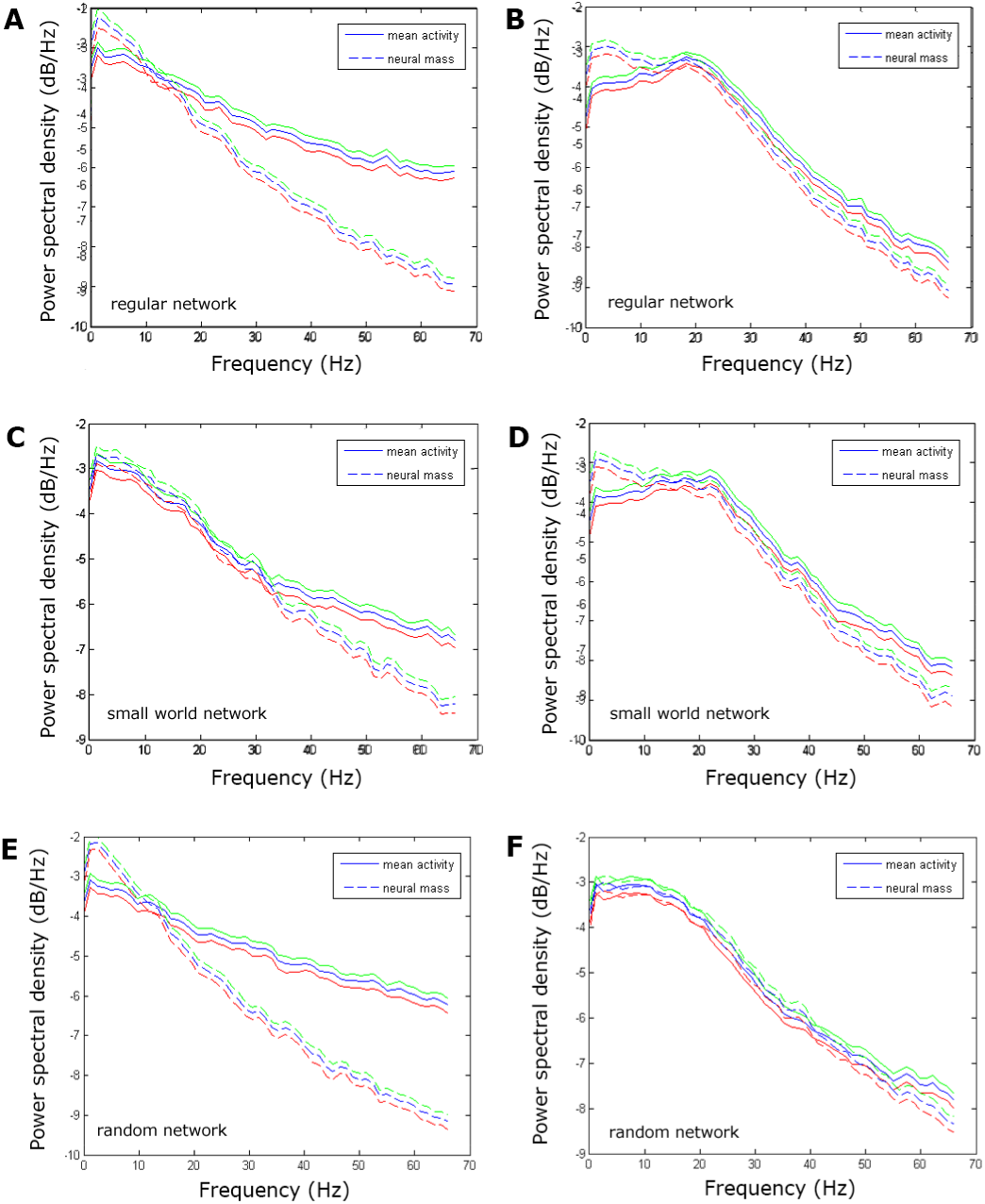
Power spectral densities. Elaboration of the simulations shown in figure 2. We have plotted the power spectral densities with confidence intervals (green and red lines) of the average network signal (normal lines) and neural mass model 1 (dashed lines) before (ACE) and after the critical thresholds (BDF) (see Figure 2 for explanation). This was done for each network configuration, regular network (AB), small-world network (CD) and a random network (DF). Note that before the threshold there is mostly overlap found for the alpha band range, except for the small world network, where this extents to the beta band. After the threshold there is a large overlap between the average network signal and the neural mass model.

## 6 Discussion

In the present study we aimed to investigate whether the average network signal of a neuronal poulation can be described by a neural mass signal when the underlying neuronal population is inhomogeneous. Our main findings are: (1) for a fully connected neuronal network there is a threshold for network size above which the neural mass model is capable of describing the average network signal; (2) By decreasing the connection density in the neuronal network we observed that for any topological configuration, there is a critical density below which there is a discrepancy between the average network signal and the neural mass signal accompanied with lower PLV values between neurons in the neuronal network; (3) this change in behaviour of the average network signals was abrupt.

Firstly, we found that there is threshold for network size, 300 neurons, above which the neural mass model is still a good description of the average network signal. We reproduced this finding with both neural mass models. These present findings are in accordance with the concept of a mean-field description, which is only valid when the number of elements in a system becomes large. However, even for large network sizes, the neural mass model has difficulty in capturing higher frequency components of the average network signal. When the number of neurons in the fully connected neuronal network increased, the firing rate also increased, leading to higher frequency components in the average network signal. However, fully connected neuronal networks are often used in computational studies (Borgers et al. 2005; Deco et al. 2013; Nakagawa et al. 2013), but do not resemble realistic neurobiological networks often characterized by sparsity of connections.

In our second analyses we found that if connections in neuronal network become too sparse there was discrepancy between the average network signal and the neural mass model. There was a critical density for all topological configurations, albeit these densities differed for the different topological configurations as for random network topology this density was around 5% in contrast to regular networks where this was 90% for a given network size. As this discrepancy between the average network signal and neural mass around this critical density coincided with a drop in PLV, we interpret these findings in the following way: synchronization of neurons in a neuronal network is a prerequisite for a valid mean-field description by a neural mass model. In all our simulations we obtained significantly different power spectral densities when we also found low PLV values. This can be understood by the notion that a decrease of synchronization between neurons in the neuronal network leads to a loss of simultaneous firing in the neuronal network and the influence of noise for the behavior of individual neurons becomes relatively larger. This leads to higher contribution of higher frequency components in the average network signal which cannot be captured by the neural mass model. Our interpretation for a requirement of a minimum amount of synchronization also explains why the density threshold for random networks was much lower than for regular networks since it is well-known that random networks are much easier to synchronize than regular networks (Barahona and Pecora 2002). Our hypothesis was further underpinned by the finding that for a larger random network the critical density was a few percent lower. This can be explained by the fact that synchronizability of a network scales with network size (Belykh et al. 2005).

The neural mass models had especially difficulty in describing high frequency components of the average network signal, especially when there was a drop in synchronization between neurons in the neuronal network. This can be understood by the notion that neural mass models can be considered as low-pass filters whose qualities are determined by the time constants (*τ* and *τ_syn_*).

Changing these parameters will lead to a change in slope of the power density spectra of the neural mass models. However, the range for which these time constants make physiologically sense is limited and therefore this restricts possible parameter values.

In the present study we used two neural mass models derived from the same LIF model, but based on different assumptions. In the first neural mass model we assumed that time scales of the membrane potential and synaptic activity were of the same order and that driving currents 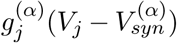 could be considered constant. This is based on the idea that fluctuations of the membrane potential around a reference value are small (Rodrigues et al. 2010). In the second neural mass model we questioned the assumption that driving currents should be considered constant and instead assumed that the time scale of the membrane potential was smaller than the time scale of synaptic activity. This led to an extra force term in Eq. 5. During our simulations we observed that the behaviour of both neural mass models was highly similar when we compared them to the average network signals, probably indicating that fluctuations of this extra force in the second neural mass model were negligible. Another assumption in both neural mass models was that communication between neurons was instantaneous and we ignored delays involved in action potential propagation and in the time course of neurotransmitter transport in synaptic clefts. The impact of delays was beyond the scope of the present paper and needs to be analysed in future studies.

The present study has implications for future model studies that try to elucidate biophysical mechanisms of disturbed oscillatory activity in neurodegenerative disorders. If the system of interest is large enough and not too sparse one can use neural mass models without concern that the neural mass model will not be a good description of the mean-field. However, when one is modelling relatively small networks and there is an indication of loss of connections between neurons, one should first check if the neurons in the underlying neuronal network are synchronized. An exception for this rule is when one is interested in alpha band activity, as we demonstrated that neural mass models were robust for this frequency range. Importantly, neural masses are generally considered to simulate activity from cortical columns. A cortical column consists of approximately 60.000-90.000 neurons/*mm*^^3^^ with a connection density of 11.05 *·* 10^^8^^*/mm*^^3^^ (Huttenlocher 1979; Boucsein et al. 2011). Note that this connection density may be around the critical threshold for networks with random network topology.

To conclude, we have used two mappings between a LIF-neuron and reduced neural mass models. Average network signals based on activity of LIF-networks can be described by neural mass models if the neurons in the network synchronize. Caution is needed when modelling neurodegenerative diseases since it is not a priori certain that synchrony is unaltered by the disease.

## 7 Appendix

### 7.1 Appendix A: A network of leaky integrate-and-fire neurons

We consider a population of *N* LIF neurons. Every neuron is described in terms of the dynamics of its membrane potential *V*_*j*_ with *j* = 1*,…, N* and with time-dependent synaptic conductances. The latter alter due to incoming spikes. Conductances are denoted as *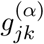*, i.e. every neuron *j* can be connected up to *N* other neurons by its synapses *k* with *k* = 1*,…, N*. We discriminate the type of synapse by superscript *α*, e.g. for excitatory and inhibitory synapses we use *α ∈* {*E, I*}. We assume that *V*_*j*_ obeys the stochastic dynamics

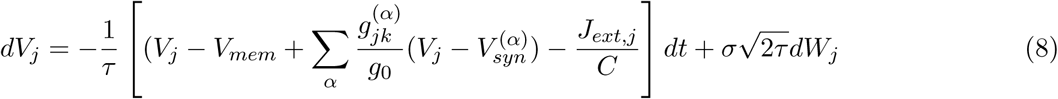

with fixed resting state potential *Vmem* and reversal potentials 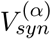 *g*_0_is the leak conductance that is considered constant and *τ* is the membrane time constant. Both constants are assumed to be identical for all neurons. The membrane time constant *τ* is related by 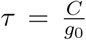 to the capacitor and membrane conductance constant *g*_0_*τ* and 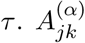 is the adjacency matrix describing whether neuron *k* targets neuron *j*

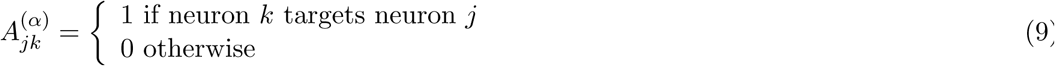

*J*_*ext,j*_ may be some external current on neuron *j*. Every neuron j is under influence of zero-centered, *δ* correlated, Gaussian white noise *W*_*j*_with unit variance, i.e. that is 𝔼 [*W*_*j*_= 0 and 𝔼 [*W*_*j*_(*t*)*W*_*j*_(*t′*)] = *δ_jk_δ*(*t t′*) where *δ_kl_* is the Kroneckers delta function and *δ*(*t - t′*) refers to Diracs delta-function. The noise strength *σ* is considered identical for all neurons *-* see Table 1 for values of all constants (Yger et al. 2011). The corresponding conductances can be described by

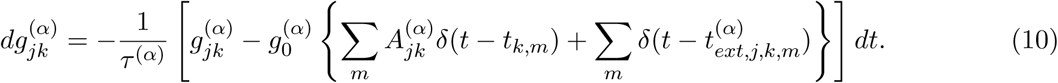

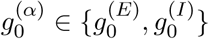 are constants related to the maximal conductance that may serve to discriminate between excitatory and inhibitory neurons. *t*_*k,m*_ is the time of the *m - th* spike of neuron *k* and, furthermore, synapse*j, k, α* may also receive external input in the form of spikes at times 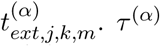 *τ* ^(*α*)^ determines the rise and decay of synaptic activity which is in general different for inhibitory and excitatory neurons. Fortunately, we can simplify Eqs. (8 & 10) when defining the total activity of the *α*-synapse of neuron *j* as the total activity of the *α*-synapse of neuron *j* as 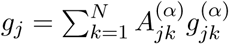 Multiplying Eq. 10 with 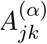 and summation over *k* yields for neuron *j* s total synapse dynamics

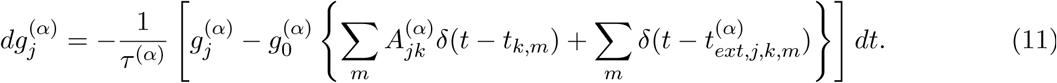

because 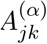 is nilpotent, i.e. 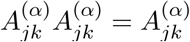 For the sake of legibility we introduce abbreviations for the incoming firing rates, namely

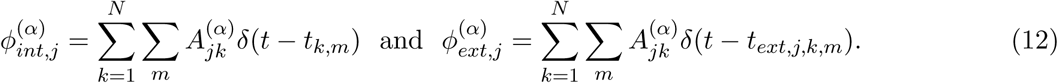

with which Eq. 11 can be rewritten as

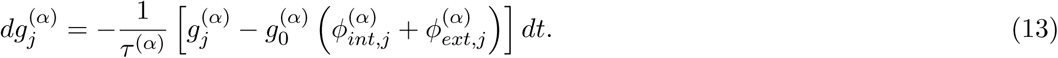

By substitution of 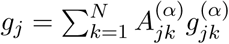 into Eq. 8, this reduces the membrane potential *V*_*j*_ into

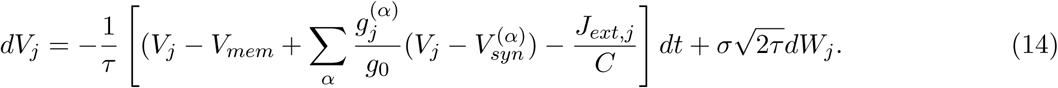

Finally, the capacitor is charged until the membrane potential *V*_*j*_ reaches a threshold, after which spikes are emitted and at which we define

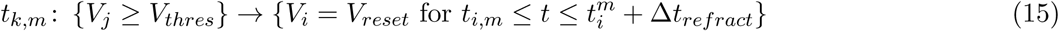

Threshold and reset potential, *V*_*thres*_ and *V*_*reset*_, respectively, as well as the refractory time Δ*t*_*refract*_, are considered identical for all neurons.

### 7.2 Appendix B: Derivation of the neural mass models

For the macro-scale we derive two neural mass models. Aim is to find the dynamics of the expectation value of the mean membrane potential

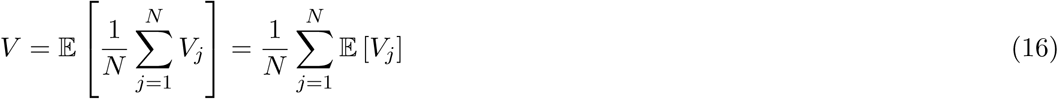

We define the expectation value of the mean synaptic conductivity as

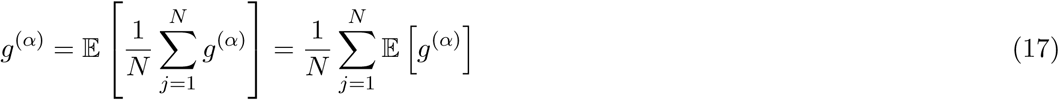

Here we assumed that 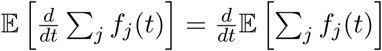 holds for all considered functions *f*_*j*_ = *V*_*j*_ and 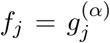 If we now combine Eqs. 14, 16 and 17 we obtain an equation describing the dynamics of the mean membrane potential *V*

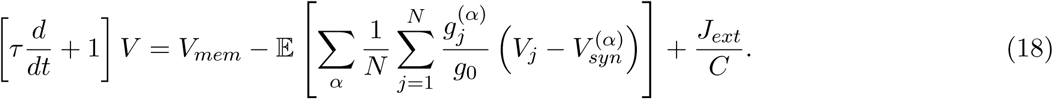

where we assumed that 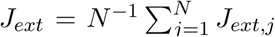 is deterministic. Similarly, we can derive the mean dynamics of the synpatic conductance if we define the expectation value of spiking behaviour

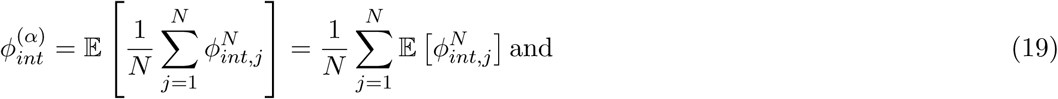

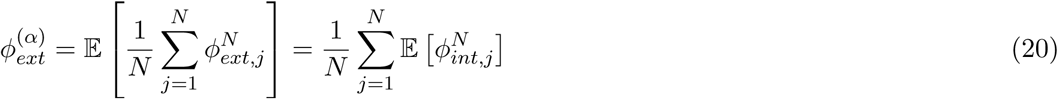

such that for we obtain an equation for the mean synaptic conductance by

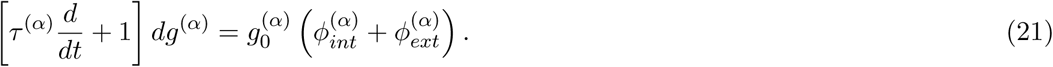

Aim of that follows is to substitute 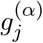 in the equation for the mean membrane potential (Eq. 18). From here we follow two separate approaches to derive neural mass models.

*Neural mass derivation 1*

In this derivation we followan approach by Rodrigues and co-workers where it is assumed that driving forces 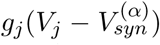 for all channels are constant (Moreno-Bote and Parga, 2005; Rodrigues et al., 2010). This is based on the assumption that fluctuations of these driving forces are small. Therefore, we ise a general, finite value for 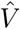 This leads to a mean-field approximation 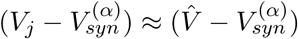 Now the mean mebrane potential’s dynamics (Eq. 18) reduces to

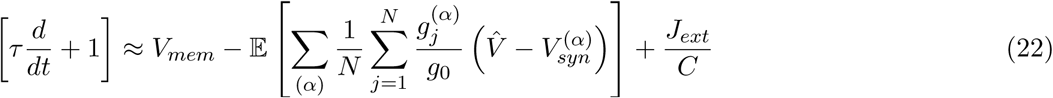

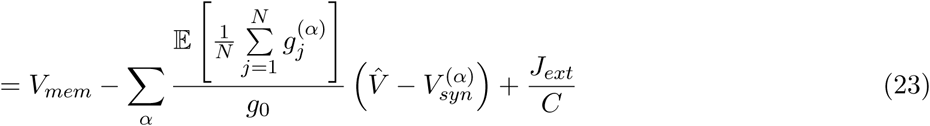

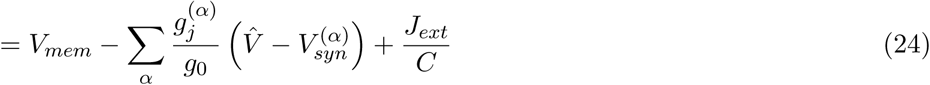

We further assume that the time scals of the mean membrane potantial and synaptic activity are of the same order irrespective of their type (Destexhe and Sejnowski, 2001; Gerstner and Kistler, 2002; Rodrigues et al., 2010). That is: *τ* ^(*α*)^ *≈ τ_syn_*. This leads to

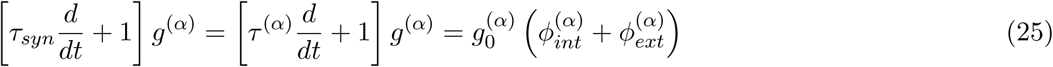

Note that the only difference between excitatory neurons/synapses is given through 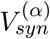 and 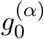 We further assumed that the external current to be constant by which 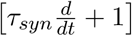 holds. Then, combining Eqs. 24 and 25 yields

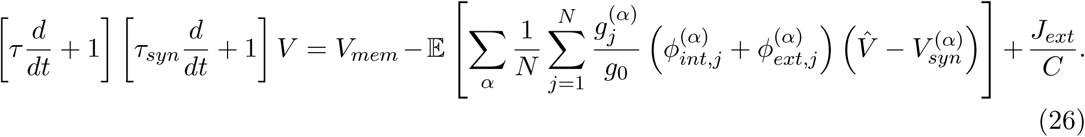

We further ignored the reset-rule and the presence of a refractory period, incorporated in the initial LIF equations, as we assume that the most important characteristics that need to be captured are sub-threshold post-synaptic potentials of neurons in the neuronal population. By inspection we find that our first neural mass model is identical to the widely known neural mass model of Freeman (Freeman, 1975)

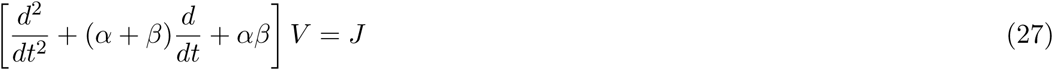

when substituting *α* = *τ-*^1^, 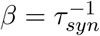 and *P* = (*ττ_syn_*)^-1^ in the RHS of Eq. 27.

*Neural mass derivation 2*

Since we are not entirely certain about the validity of all assumptions made in the first neural mass derivation, especially the assumption that the driving forces 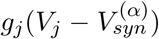 are constant, we search for an alternative derivation. Now we start again with the equation describing the dynamics of *V*, Eq. 18.

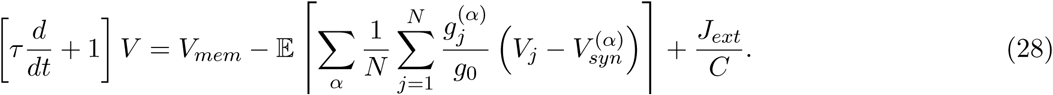

We now assume that the synaptic time scales of the excitatory and inhibitory conductances for individual neurons are again in the same order of magnitude. This indicates that we can write

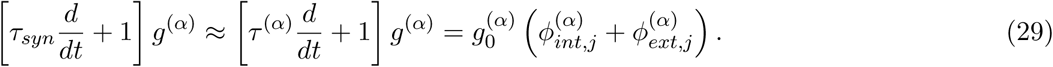

If we now multiply Eq. 28 by the LHS of Eq. 29 then we can write

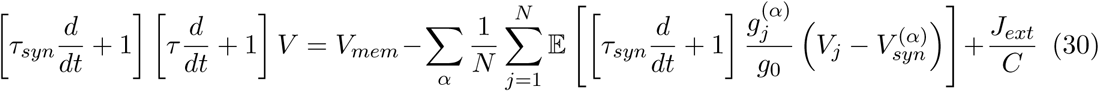

Subsequently, by using the product rule for differentiation this leads to

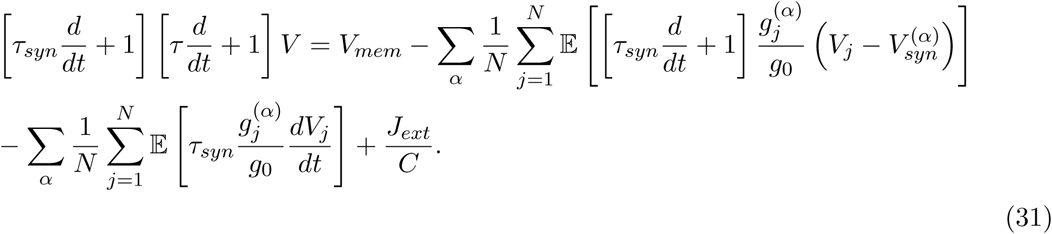

Furthermore, by substituting the RHS of Eq. 29 in Eq. 31 we obtain

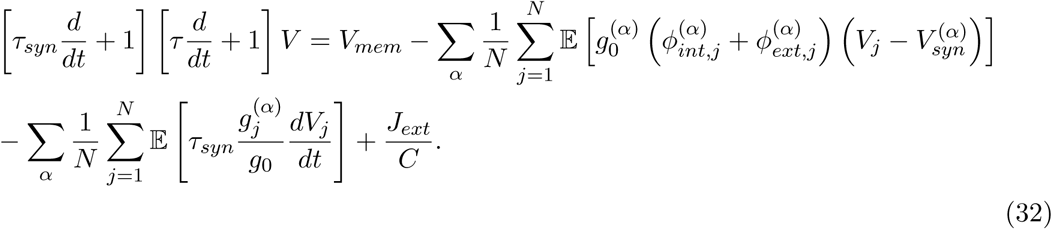

Next we assume the firing rates to be deterministic. This allows for a mean-field approximation such that 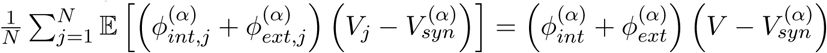 replaced in the first term of RHS of Eq. 32. To treat the final term of Eq. 32 we compare the time scale of the membrane potential vis-a-vis with the synaptic conductivity. If the first is considered to be much smaller than the latter, meaning that the membrane potential instantly follows changes at the synapse, the dynamics of the membrane potential can be eliminated adiabatically that might be formalized as

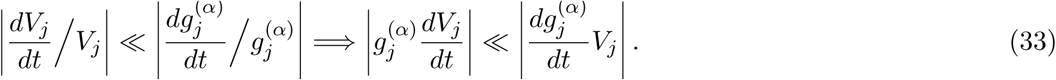

This allows for reducing Eq. 32 to

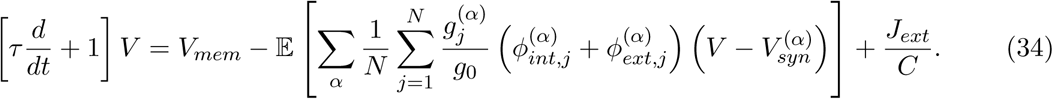

which gives us the equation for the second neural mass model. Qualitative difference to Eq. 26 is that Eq. 34 contains an extra parametric forcing term 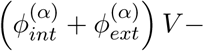 which does no longer agree with Freemans model.

### 7.3 Appendix C: Algorithms used in simulations

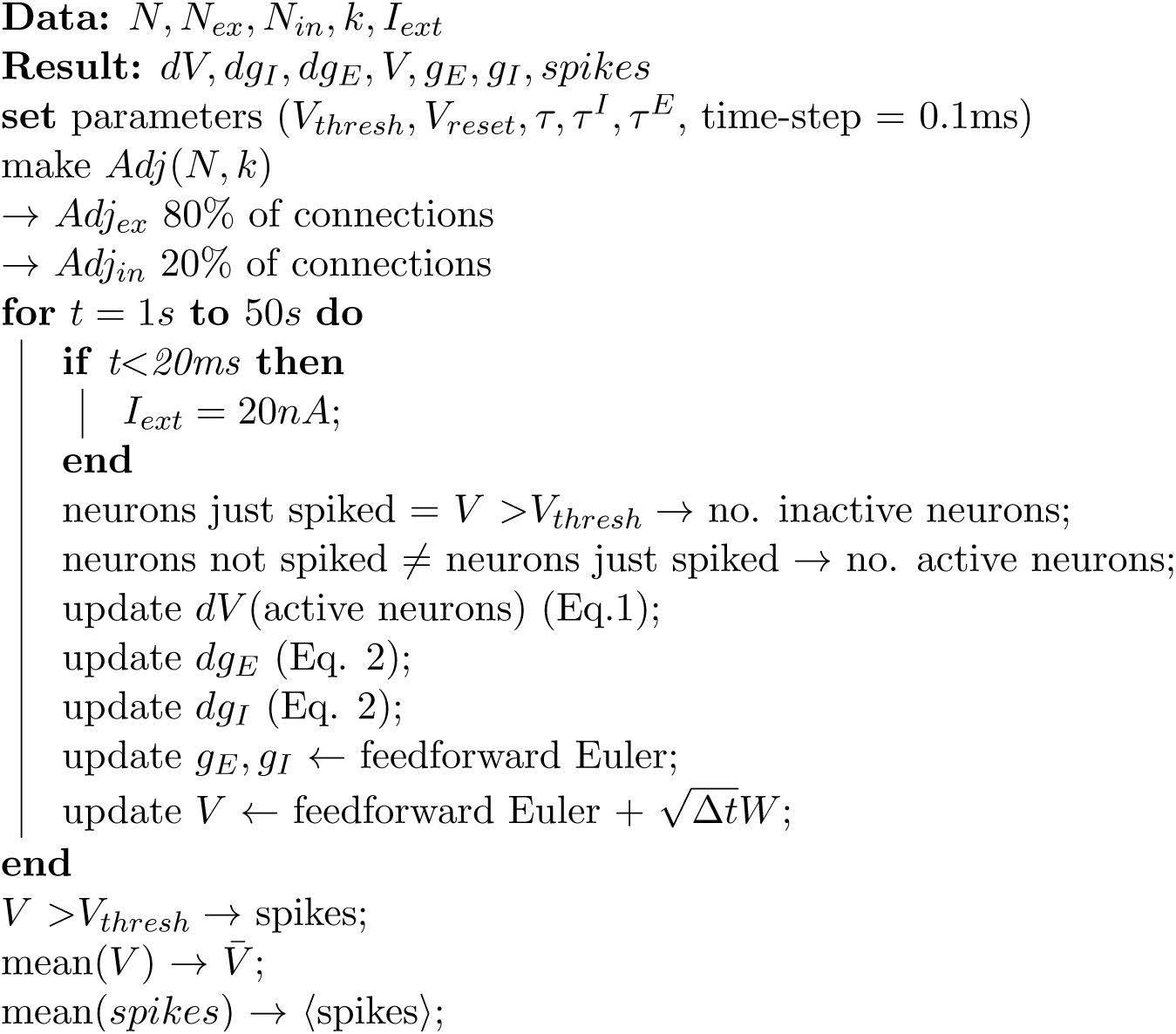

**Algorithm 1:** Simulation of a LIF network

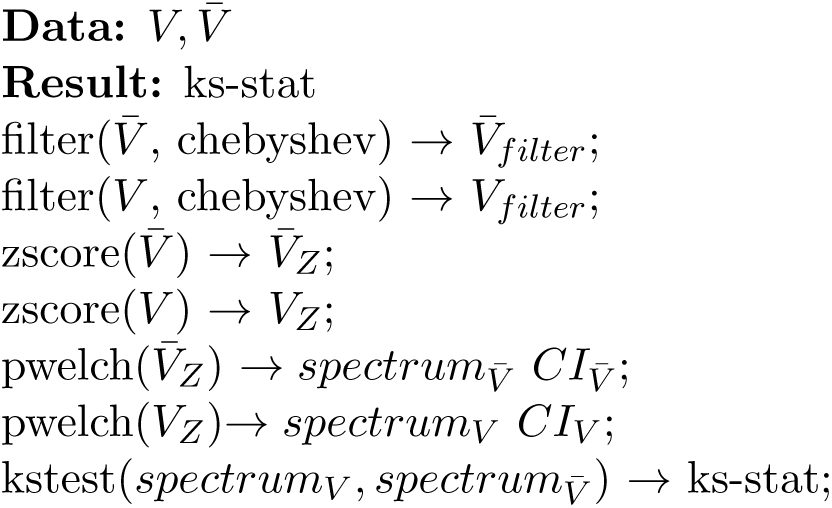

**Algorithm 2:** Comparison of spectra between average network signals (V) and neural mass signals *V*

## References

Abbott, L., and van Vreeswijk, C. (1993). Asynchronous states in networks of pulse-coupled neuron. Phys. Rev. A 48, 1483–1490.

Amari, S.I. (1972). Characteristics of Random Nets of Analog Neuron-Like Elements. IEEE Trans. Syst. Man Cybern 2, 643–657.

Amari, S.-I., Yoshida, K., and Kanatani, K.-I. (1977). A mathematical foundation for statistical neurodynamics. SIAMJ. Appl. Math 33, 95–126.

Baladron, J., Fasoli, D., Faugeras, O., and Touboul J. (2012). Mean-field description and propagation of chaos in networks of Hodgkin-Huxley and FitzHugh-Nagumo neurons. J Math Neurosci 2, 2190–8567.

Barahona, M., and Pecora, L. (2002). Synchronization in Small-World Systems. Phys. Rev. Lett. 89, 054101–1-054101-4.

Belykh, I., de, L.E., and Hasler, M. (2005). Synchronization of bursting neurons: what matters in the network topology. Phys. Rev. Lett. 94, 188101.

Borgers, C., Epstein, S., and Kopell, N.J. (2005). Background gamma rhythmicity and attention in cortical local circuits: a computational study. Proc. Natl. Acad. Sci. U. S. A 102, 7002–7007.

Boucsein, C., Nawrot, M.P., Schnepel, P., Aertsen, A. (2011). Beyond the cortical column: abundance and physiology of horizontal connections imply a strong role for inputs from the surround. front. in neuroscience 5. 1-13.

Breakspear, M., Terry, J.R., and Friston, K.J. (2003). Modulation of excitatory synaptic coupling facilitates synchronization and complex dynamics in a biophysical model of neuronal dynamics. Network. 14, 703–732.

Brunel, N., and Hakim, V. (1999). Fast global oscillations in networks of integrate-and-fire neurons with low firing rates. Neural Comput 11, 1621–1671.

Cessac, B. (1995). Increase in complexity in random neural networks. J. Phys. I (France) 5, 409–432.

Cessac, B. (2008). A discrete time neural network model with spiking neurons: rigorous results on the spontaneous dynamics. J. Math. Biol. 56, 311–345.

Cessac, B., and Vieville T (2013). On dynamics of integrate-and-fire neural networks with adaptive conductance based synapses. Front. Neurosci 2.

Creutzfeldt, O., and Ito, M. (1968). Functional synaptic organization of primary visual cortex neurones in the cat. Exp. Brain Res. 6, 324–352.

Deco, G., Jirsa, V., Robinson, P., Breakspear, M., and Friston, K. (2008). The dynamic brain: from spiking neurons to neural masses and cortical fields. Plos Comput Biol 4.

Deco, G., Ponce-Alvarez, A., Mantini, D., Romani, G.L., Hagmann, P., and Corbetta, M. (2013). Resting-State Functional Connectivity Emerges from Structurally and Dynamically Shaped Slow Linear Fluctuations. J. Neurosci. 33, 11239–11252.

Destexhe, A., and Sejnowski, T.J. (2001). Thalamocortical Assemblies: How Ion Channels, Single Neurons and Large-Scale Networks Organize Sleep Oscillations. Oxford University Press).

Erdos, P., Renyi, A. (1960). On the evolution of random graphs. Magyar Tud. Akad. Mat. Kutato Int. Kozl, 5, 17–61.

Faugeras, O., Touboul, J., and Cessac, B. (2009). A constructive mean-field analysis of multipopulation neural networks with random synaptic weights and stochastic inputs. front. in comp. neuroscience 3, 1–28.

Freeman, W. (1975). Mass action in the nervous system.

Gerstner, W., and Kistler, W. (2002). Spiking neuron models: single neurons, populations, plasticity. Cambridge University Press, Cambridge).

Grabska-Barwinska, A., and Latham, P.E. (2013). How well do mean field theories of spiking quadratic-integrate-and-fire networks work in realistic parameter regimes? J. Comput. Neurosci.

Huttenlocher, P.R. (1979). Synaptic density in human frontal cortex developmental changes and effects of aging. Brain Reseacrch 163, 195–205.

Landau, L.D., and Lifshitz, E.M. (1968). Statistical Physics. In Course in Theoretical Physics, pp. 1–28.

Mardia, K.V. (1972). Statistics of directional data. London: academic press.

Mattia, M., and Del Giduice, P. (2002). Population dynamics of interacting spiking neurons. Phys. Rev. E Stat. Nonlin. Soft Matter Phys. 66, 51917.1-51917.19.

Molgedey, L., Schuchardt, J., and SChuster, H. (2013). Suppressing chaos in neural networks by noise. Phys. Rev. Lett. 69, 3717–3719.

Moreno-Bote, R., and Parga, N. (2005). Simple model neurons with AMPA and NMDA filters: role of synaptic time scales. Neurocomputing 65-66, 441–448.

Moynot, O., and Samuelides, M. (2002). Large deviations and mean-field theory for asymmetric random recurrent neural networks. Probab. Theory Relat. Fields 123, 41–75.

Nakagawa, T.T., Woolrich, M., Luckhoo, H., Joensson, M., Mohseni, H., Kringelbach, M.L., Jirsa, V., and Deco, G. (2013). How delays matter in an oscillatory whole-brain spiking-neuron network model for MEG alpha-rhythms at rest. Neuroimage.

Robinson, P., and Kim, J. (2010). Spike, rate, field, and hybrid methods for treating neuronal dynamics and interactions. Journal of Neuroscience Methods 205, 283–294.

Rodrigues, S., Chizhov, A., Marten, F., and terry, J. (2010). Mappings between a macroscopic neural-mass model and a reduced conductance-based model. Biol. Cybern. 361-371.

Rudolph-Lilith, M., Dubois, M., and Destexhe, A. (2012). Analytical integrate-and-fire neuron models with conductance-based dynamics and realistic postsynaptic potential time course for event-driven simulation strategies. Neural Comput. 24, 1426–1461.

Samuelides, m., and Cessac, B. (2007). Random recurrent neural networks. Eur. Phys. J. Spec. Top 142, 7–88.

Touboul, J.D., and Ermentrout, G.B. (2011). Finite-size and correlation-induced effects in meanfield dynamics. J. Comput. Neurosci. 31, 453–484.

Treves, A. (1993). Mean-field analysis of neuronal spike dynamics. Netw. Comput. Neural Syst 4, 259–284.

Vogels, T.P., and Abbott, L.F. (2005). Signal propagation and logic gating in networks of integrate-and-fire neurons. J. Neurosci. 25, 10786–10795.

Watts, D.J., Strogatz, S.H. (1998). Collective dynamics of ‘small-world’ networks. Nature 393, 440–442.

Wilson, M., Robinson, P., O’Neill, B., and Steyn-Ross, D. (2012). Complementarity of Spike- and Rate-Based Dynamics of Neural Systems. Plos Comput Biol 8, 1–19.

Wong, K.F., and Wang, X.J. (2006). A recurrent network mechanism of time integration in perceptual decisions. J. Neurosci. 26, 1314–1328.

Yger, P., El, B.S., Destexhe, A., and Fregnac, Y. (2011). Topologically invariant macroscopic statistics in balanced networks of conductance-based integrate-and-fire neurons. J. Comput. Neurosci. 31, 229–245.

